# An anisotropic attraction model for the diversity and robustness of cell arrangement in nematodes

**DOI:** 10.1101/132654

**Authors:** Kazunori Yamamoto, Akatsuki Kimura

## Abstract

In early embryogenesis stages in animals, cells are arranged into a species-specific pattern in a robust manner. The cell arrangement patterns are diverse, even among close relatives. We evaluated how the diversity and robustness are achieved in developing embryos. We succeeded in reproducing different patterns of cell arrangements observed in various nematode species using *Caenorhabditis elegans* embryos by changing the eggshell shapes. This implies that the diversity of cell arrangements can be explained by differences in a shape parameter. Additionally, we found that the cell arrangement was robust against eggshell deformation. Computational modeling revealed that, in addition to repulsion forces, attraction forces are sufficient for this robustness. Genetic perturbation experiments demonstrated that attraction forces derived from cell adhesion are necessary for the robustness. The proposed model accounts for both diversity and robustness of cell arrangements and contributes to our understanding of how diversity and robustness are achieved in developing embryos.

## Introduction

In multicellular organisms, nearby cells communicate with each other by sending and receiving signals mediated by their surface molecules (Alberts et al., 2008). The cell arrangement pattern, which refers to the pattern of cell-cell contacts, is important for development and homeostasis of an organism. During embryogenesis, specific cell arrangement patterns are established and specific cell-cell contacts in the patterns define cell fate and body plan (Gilbert, 2016). Although the importance of the cell arrangement pattern is well studied, the mechanisms of how specific arrangements are established are not fully understood.

In this study, we investigated the mechanical basis underlying the diversity and robustness of cell arrangement patterns using four-cell stage nematode embryos as models. Cell arrangement patterns at the four-cell stage are diverse among nematode species (Goldstein, 2001). The diversity of cell arrangement patterns is often explained by a variation in the orientation and the position of the mitotic spindle (Akiyama et al., 2010; Pierre et al., 2016), and in the four-cell-stage nematode embryos in particular (Schulze and Schierenberg, 2011). However, the mitotic spindle is not the sole determinant of the cell arrangement. Cells move and change their arrangement after cell division depending on their interactions with other cells in a confined space. In the nematode embryo, this confined space is defined by the eggshell (Olson et al., 2012). At the four-cell stage, the *Caenorhabditis elegans* embryo generally takes a “diamond” type of cell arrangement (Fig. 1A) inside the eggshell, and a “T-shaped” type (Fig. 1A) when the eggshell is removed (Edgar et al., 1994). Interestingly, the eggshell shapes, too, are diverse among nematode species (Goldstein, 2001). We noticed that there seems to be a correlation between eggshell shapes and cell arrangement patterns. We thus hypothesized that the diverse patterns of cell arrangements are produced by the diverse shapes of the eggshells. Because the effect of eggshell shape on the pattern of cell arrangement has not been examined, we attempted to change the shapes of the *C. elegans* eggshells to assess whether it is a source of diversity in cell arrangement patterns.

**Figure 1.**
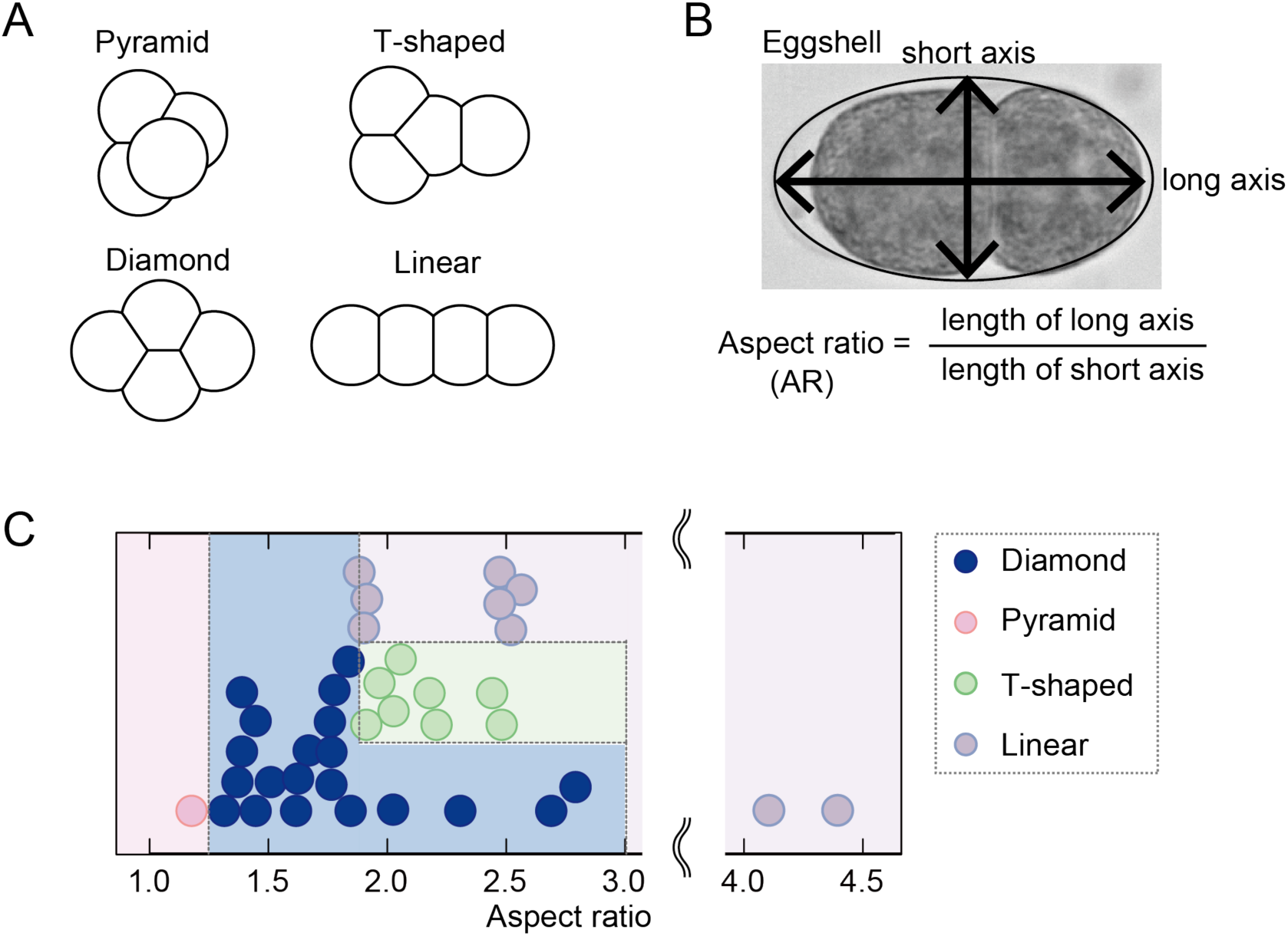
Cell arrangement patterns in various nematode species. (A) Classification of the cell arrangement patterns. Depending on the cell–cell contacts, the patterns at the four-cell stage are classified into “pyramid,” “diamond,” “T-shaped,” or “linear” types. (B) The aspect ratio (AR) was calculated as the long axis length divided by the short axis length of the eggshell. (C) Relationship between the cell arrangement pattern and the AR in embryos of various nematode species. All data are summarized in Table S1.

While cell patterns are diverse, individual species often take a specific pattern reproducibly in nematode species (Gilbert, 2016; Schulze and Schierenberg, 2011). Such a robust pattern is critical for embryo development. *C. elegans* is one of the organisms for which specific cell-cell contacts and their roles in development are well studied. At the four-cell stage, the blastomeres take the diamond-type arrangement in which two pairs of cells (EMS and P2, or ABp and P2 cells) contact (Fig. S1A) and send signals mediated by Wnt-Frizzled or Notch-Delta, both of which are critical to establish the dorsal-ventral embryo axis (Gönczy and Rose, 2005). Changes in the arrangement pattern have deleterious effects on the embryo (Goldstein, 1992; Kemphues et al., 1988). To date, the robustness of cell arrangement has not been examined in a systematic manner. In this study, we examined how robust the diamond-type arrangement is against deformation of the eggshell using *C. elegans* embryos.

Mechanistic bases for the diversity and robustness of cell arrangements can be understood by constructing theoretical models. A good mechanical model that accounts for the diamond-type of cell arrangement was reported previously (Fickentscher et al., 2013). The model assumes two types of repulsion forces: a repulsion force between cells, and a repulsion force between a cell and the eggshell. The model succeeded to reproduce both position and trajectory of cells precisely up to the 12-cell stage for embryos with an average shape (Fickentscher et al., 2013). Repulsion forces are commonly assumed to explain the patterns of cell arrangements in various species (Zammataro et al., 2007; Akiyama et al., 2010; Kajita et al., 2003). Such repulsion forces are provided by the surface tension and elasticity of the cell (Fletcher et al., 2014; Fujita and Onami, 2012). However, it has not been examined whether the model based on the repulsion forces also accounts for the diversity and robustness of cell arrangements.

In this study, we focused on embryo deformation as a mechanical perturbation to investigate the diversity and robustness. The purposes of this study were (i) to test whether the shape of the eggshell accounts for the diversity of the cell arrangement patterns in four-cell nematode embryos, (ii) to characterize the robustness of the diamond pattern of *C. elegans* against deformation, (iii) to construct a theoretical model to account for the diversity and robustness, and (iv) to understand the molecular basis of the model.

## Results

### Eggshell shape and cell arrangement pattern are correlated in various nematode species

To examine whether the eggshell shape is related to the diversity of cell arrangements at the four-cell stage, we investigated the correlation between eggshell shapes and cell arrangement patterns in various nematode species. Based on images in published reports (Goldstein, 2001; Schulze and Schierenberg, 2011), the patterns of cell arrangements at four-cell stage were classified into “diamond,” “pyramid,” “T-shaped”, or “linear” types, which are defined by cell–cell contacts (Fig. 1A). We quantified the eggshell shape on the basis of the aspect ratio (AR), which is calculated by dividing the length of the long axis by that of the short axis of the eggshell (Fig. 1B), and associated them with the pattern of cell arrangement (Table S1). For *Diploscapter coronata* (Lahl et al., 2009) and *Aphelenchoides besseyi* (Yoshida et al., 2009), we obtained animals and imaged the embryos ourselves (Table S1). We then examined the relationship between the ARs and the cell arrangement patterns (Fig. 1C). The patterns seemed to correlate with the ARs, supporting the notion that the diversity in eggshell shapes can be a source of the diversity in cell arrangement patterns. The pyramid-type of cell arrangement was observed at low ARs (AR = 1.2). The diamond-type was observed 100% at ARs from 1.4 to 1.8. For ARs over 4.0, only the linear type was observed.

Shape is not a sole determinant of cell arrangement because different patterns were observed for similar AR values. For ARs from 1.9 to 2.8, the diamond, T-shaped, and linear types were observed, depending on the species (Fig. 1C). The diversity independent of the shapes is likely caused by a diversity in the orientations of cell division (Schulze and Schierenberg, 2011). For example, in embryos taking the linear-type arrangement at the four-cell stage, the cells at the two-cell stage both divide along the long axis of the egg. However, we noticed that, in a species taking the linear type (*Zeldia*), the cells at the two-cell stage likely divide perpendicular to each other (Schulze and Schierenberg, 2011). Because the species has a slender eggshell (AR = 2.5, Table S1), we suspected that the eggshell shape induced the linear-type arrangement. In summary, we suspected that the shape of eggshell is a major parameter determining the pattern of cell arrangement and can generate diversity therein.

### Deformation of eggshell shape in *C. elegans* mutants and RNAi-treated strains

To test whether deformation of the eggshell shape can change the cell arrangement pattern, we searched for genes that affect eggshell shape in *C. elegans*. In the WormBase database (<u>www.wormbase.org</u>), seven genes were categorized as “egg shape morphology variant.” Among these genes, four were excluded because their knockdown phenotype was embryonic lethal. The ARs of mutant strains of two genes (*spe-9*, *ceh-18*) were not significantly different from those of the wild types in our analyses (Fig. S1B). Thus, the only remaining gene was *lon-1*. The ARs of *lon-1*(*e185*) mutant embryos (1.8 ± 0.2, *n* = 258) were significantly larger than those of the wild type (1.6 ± 0.1, *n* = 281) (Fig. 2A). A previous study in our laboratory (Hara and Kimura, 2009) had demonstrated that RNAi-mediated knockdown of *C27D9.1* might increase the AR. Here, we investigated this possibility further. The results demonstrated that RNAi of this gene indeed increased the AR (1.9 ± 0.2, *n* = 90) (Fig. S1B). Furthermore, we succeeded in obtaining high-AR embryos by knocking down *C27D9.1* by RNAi with *lon-1*(*e185*) mutant background (2.3 ± 0.3, *n* = (Fig. 2A).

**Figure 2.**
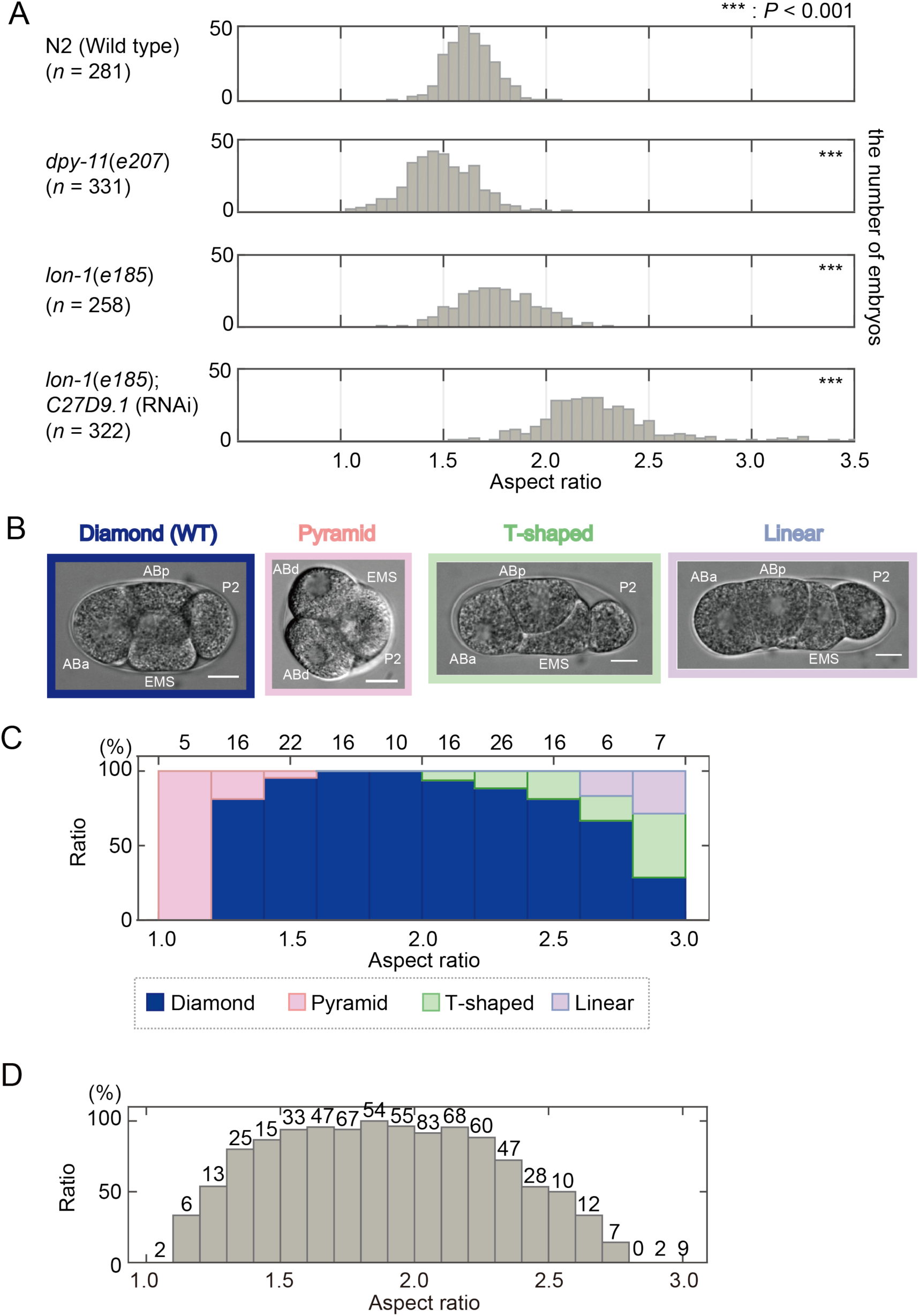
Eggshell shape determines cell arrangement pattern in the *C. elegans* embryo. (A) Histograms show the ARs in *C. elegans* mutants and RNAi-treated strains. The means ± SDs of the eggshell shapes in each strain are as follows: N2 (wild type) (1.6 ± 0.1, *n* = 281), *dpy-11*(*e207*) (1.5 ± 0.2, *n* = 331), *lon-1*(*e185*) (1.8 ± 0.2, *n* = 258), *lon-1*(*e185*); *C27D9.1* (RNAi) (2.3 ± 0.3, *n* = 322). For statistical analysis, normality was checked by a Shapiro–Wilk test, and homoscedasticity was confirmed using an F test. \*\*\**P* < 0.001 vs. N2 (wild type). Student’s *t*-test was used for *dpy-11*(*e207*), *lon-1*(*e185*). Wilcoxon’s rank sum test was used for *lon-1*(*e185*); *C27D9.1* (RNAi). (B) Micrographs showing the different cell arrangement patterns at the four-cell stage of *C. elegans* embryos. For the pyramid-type arrangement, the daughter cells of the AB cell were marked as ‘ABd‘ because we cannot distinguish ABa and ABp in this arrangement. Scale bars are 10 μm. (C) Relationship between rate of appearance of the four types of cell arrangement (blue, diamond type; pink, pyramid type; green, T-shaped type; purple, linear type) and the ARs (*n* = 188). The data include four strains: N2, *dpy-11*(*e207*), *lon-1*(*e185*), and *C27D9.1* RNAi-treated strains were on *lon-1*(*e185*) background. The numbers above the bars represent the number of embryos. (D) Dependence of hatch rate on AR (*n* = 643). The data include five strains: N2, *dpy-11*(*e207*), *lon-1*(*e185*), and *C27D9.1* RNAi-treated strains were on N2 or *lon-1*(*e185*) background. The numbers above the bars represent the number of embryos.

Next, we attempted to obtain embryos with the ARs lower than those of the wild type. Because we obtained long (high-AR) embryos from the *lon-1* mutant strain, whose adult body shape is also long (Maduzia et al., 2002; Morita et al., 2002), we speculated that short (low-AR) embryos might be obtained from short adults (Fig. S1C). We examined a mutant strain with short adults, *dpy-11*(*e207*) (Ko and Chow, 2002), and found that it produced low-AR embryos (1.5 ± 0.2, *n* = 331) (Fig. 2A).

In a previous study, an *spv-1* mutant was reported to produce embryos of various shapes (Tan and Zaidel-Bar, 2015) (Fig. S1B). We did not use the *spv-1* mutant in the present study because it produces embryos with non-ellipsoidal shape and/or with very small or very large volumes. In summary, we were able to obtain *C. elegans* embryos with ARs ranging from 1.0 to 3.5. The genes used to deform the eggshell shape (i.e. *lon-1*, *C27D9.1*, and *dpy-11*) are non-essential genes and did not affect the initial orientation of the mitotic spindle up to the four-cell stage (Fig. S2A).

### Patterns of cell arrangements are changed in eggshell shape variants of the *C. elegans* embryos

To evaluate the effect of eggshell shape on the pattern of cell arrangement, we observed cell arrangements at the four-cell stage in different eggshell shapes. Normally, at the four-cell stage, *C. elegans* embryos take the diamond-type of cell arrangement, in which the nuclei are positioned at the vertexes of a diamond shape (Fig. S1A). The embryonic cells of *C. elegans* are named after the mother cell and depending on their position relative to the sister cells (Sulston et al., 1983). At the four-cell stage, all cells (ABa, ABp, EMS, and P2) except ABa and P2 make contact with each other (Fig. 2B, S1A).

We succeeded in changing cell arrangements by changing the eggshell shapes (Fig. 2B). When the AR decreased below 1.5 (i.e., the eggshell becomes spherical), the pyramid-type of cell arrangement was observed, in which all four cells, including ABa and P2, made contact (Fig. 2B,S2B). The pyramid-type of cell arrangement was dominant when the AR was below 1.2 (Fig. 2C). In contrast, when the AR exceeded 2.0, the T-shaped type of cell arrangement appeared, in which neither ABp and P2 nor ABa and P2 made contact (Fig. 2B, S2B). The T-shaped type was dominant when the AR exceeded 2.8 (Fig. 2C). When the AR exceeded 2.7, the linear type appeared, in which the four nuclei are arranged linearly and ABa and EMS, ABa and P2, and ABp and P2 do not contact (Fig. 2B, 2C, S2B). These changes in the pattern of cell arrangement should not be the direct consequence of mutation or knockdown of the targeted genes because (i) the patterns varied even among embryos of the same genotype in an AR-dependent manner (Fig. S2B), and (ii) the gene manipulations used did not alter the initial direction of cell division, in which ABa and ABp divide perpendicular to the anterior-posterior axis, whereas EMS and P2 divide parallel to the axis (Fig. S1A, S2A).

Overall, these results demonstrated that the pattern of cell arrangement can be changed by changing the shape (i.e., the AR) of the eggshell. The hypothesis that eggshell shape contributes to diversity in cell arrangements was supported by the fact that the different patterns of cell arrangement observed in different nematode species (Fig. 1C) could be reproduced in *C. elegans* embryos with different eggshell shapes (Fig. 2C).

### Robustness of the diamond-type cell arrangement in the *C. elegans* embryo

From the experiment on changing the eggshell shape, we noticed that the normal arrangement (i.e., the diamond type for the *C. elegans*) was dominant in a wide range of ARs (from 1.3 to 2.8) (Fig. 2C). This range includes that observed for wild-type eggshells, which vary from 1.3 to 2.0 (Fig. 2A), and is consistent with the range of ARs in other nematode species displaying the diamond-type arrangement (Fig. 1C).

To correlate the robustness of the diamond-type arrangement with the robustness of embryogenesis, we quantified the hatching rate of embryos with different eggshell shapes. The hatching rate is the proportion of embryos that hatch from the eggshell to become L1 larvae, implying normal embryogenesis. The hatching rate decreased with increasing deviance of the AR from that of the wild type (Fig. 2D, S2C), and correlated with the rate of embryos taking the diamond-type arrangement (Fig. 2C). The failure of embryogenesis at low ARs was consistent with the report of the *spv-1* mutant (Tan and Zaidel-Bar, 2015), and indicates the pyramid-type arrangement (abnormal contact between ABa and P2 cells) has a deleterious effect on embryogenesis. The failure of embryogenesis at high ARs is consistent with the notion that the contact between ABp and P2 cells is important for dorsal-ventral axis formation (Gönczy and Rose, 2005; Priess, 2005). We confirmed that no embryos developed to hatching if they took a pattern of cell arrangement other than the diamond type. Overall, the correlation between the hatching rate and the diamond type of cell arrangement supported the notion that the robustness of the diamond type at the four-cell stage is critical for the robustness of embryogenesis against eggshell deformation.

### Computer simulation of the repulsion-only (RO) model

To understand the mechanical basis of the cell arrangements, we asked whether an existing mechanical model of the cell arrangements accounts for the diversity and robustness of the cell arrangements against deformation of the eggshell. The model constructed by Fickentcher *et al*. (2013) accurately accounts for the arrangements and trajectory of cells from the two-cell to 12-cell stages of *C. elegans* embryos in case of a normal eggshell shape (i.e., AR = 1.7 (Fickentscher et al., 2013)) and under the condition that the direction, timing, and volume ratio of cell division are provided. The model considers two types of repulsion forces: the repulsion between neighboring cells and the repulsion between a cell and the eggshell (Fig. 3A). In this report, we termed this model the repulsion-only (RO) model. In this model, the strength of the repulsion force depends on the distance between the centers of two cells (Fig. 3B) or between the cell center and the nearest part of the eggshell. Here, the cell center is defined as the center of the nucleus.

**Figure 3.**
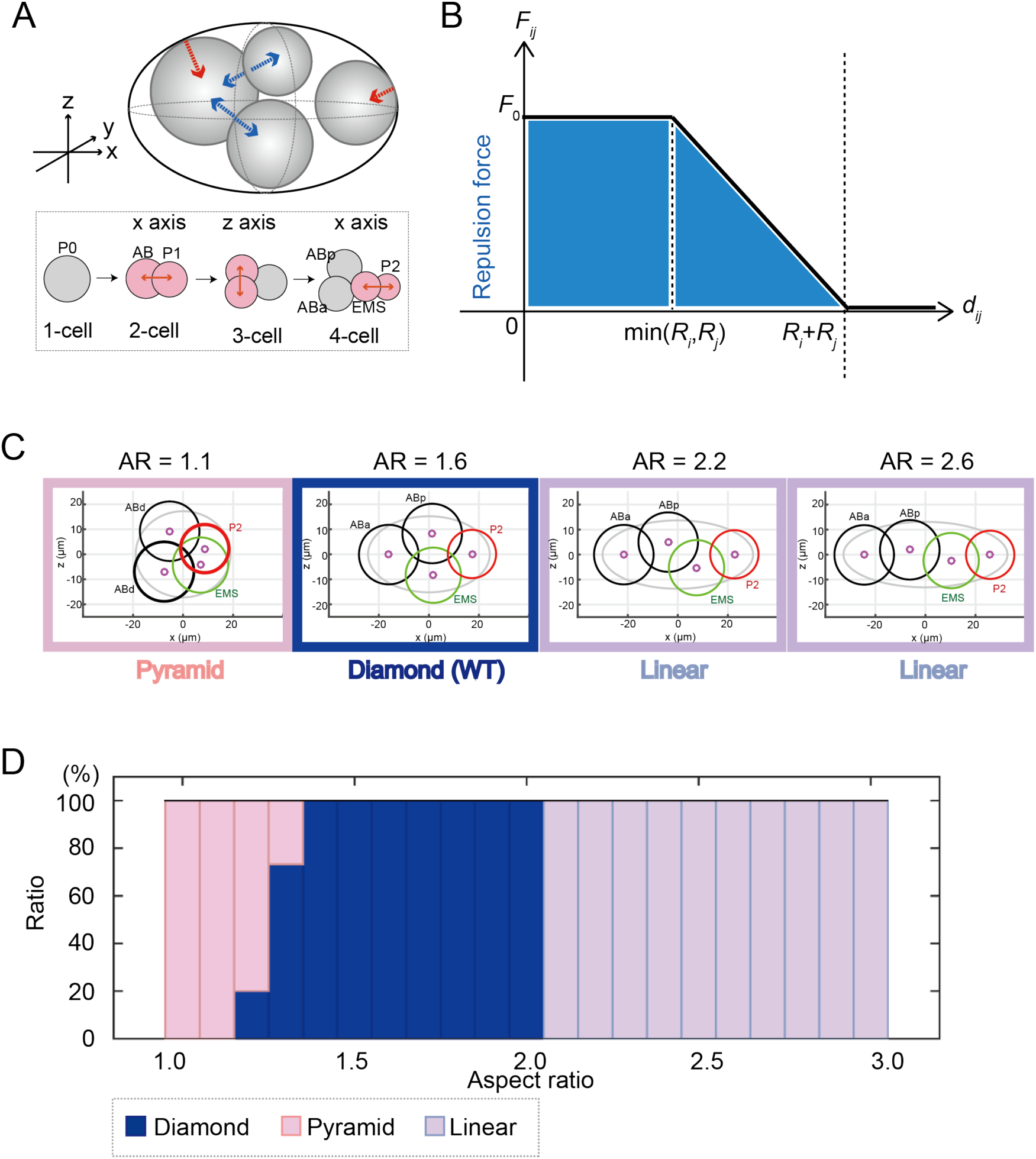
Simulation of cell arrangement patterns with the repulsion force-only (RO) model. (A) Schematic representation of the RO model. The upper panel shows the repulsion forces between the cells (blue arrows) and between the cells and the eggshell (red arrow). The lower panel shows the orientation and timing of cell divisions. (B) Relationship between strength of intercellular force (*F*_*ij*_) and cell–cell distance (*d*_*ij*_) in the RO model. (C) Examples of the cell arrangement patterns at the four-cell stage simulated by the RO model at AR = 1.1, 1.6, 2.2, and 2.6 respectively. (D) Relationship between rates of appearance of different patterns of cell arrangements and the ARs in the RO model (blue, diamond type; pink, pyramid type; purple, linear type).

We examined whether the RO model also reproduces the diverse patterns of cell arrangements when the shape of the eggshell is changed while the other parameters are maintained. The diamond-type arrangement was observed in 100% of simulations with ARs from 1.4 to 2.0 (Fig. 3C, 3D). Interestingly, when the AR exceeded 2.0, the diamond type was not observed anymore and 100% of simulations resulted in the linear-type arrangement (Fig. 3C, 3D). This did not reflect the situation *in vivo*, where the diamond type decreased gradually when the AR exceeded 2.0, and more than 50% of the embryos showed the diamond type even for ARs from 2.6 to 2.8 (Fig. 2C). Therefore, the RO model was less robust against eggshell deformation than real embryos. Another notable difference between the real embryos and the RO model was that the model did not reproduce the T-shaped arrangement at any AR (Fig. 3D), in contrast to real embryos, in which this type of arrangement was observed for ARs over 2.0 (Fig. 2C). Instead, 100% of the linear type was obtained in the RO model for ARs over 2.0 (Fig. 3D), whereas this type was observed in the real embryos only when the AR exceeded 2.7 (Fig. 2C).

Because the RO model was originally developed to reproduce the cell arrangement in a normal shape (AR = 1.7), and not for diversity or robustness, we tested whether the RO model can reproduce the diversity and robustness by changing other parameters of the model. We focused on the parameter of repulsion force strength because the other parameters were based on experimental measurements. Moreover, because the absolute values of the forces affect only division speed but not the final (stable) position of the cell, we changed the ratio of repulsion forces between cell–cell (*F*_0_) and cell–eggshell (*K*_0_). We first determined the range for cells taking the diamond-type arrangement in a normal shape. Next, within this ratio range, we changed the shape and examined whether the diversity and robustness were reproduced. The trend in frequency patterns of cell arrangements did not change largely (Fig. S3). From these results, we concluded that the RO model is not sufficient to explain the diversity and robustness of cell arrangements against eggshell deformation.

### Characterization of the intercellular forces in the *C. elegans* embryo

To unravel mechanisms for the diversity and robustness of cell arrangements upon eggshell deformation (i.e., ARs > 2.0), we carefully observed the physical interactions between embryonic cells. To characterize the nature of these physical interactions, we removed the eggshell. We noticed that, even when the cells are not confined inside the eggshell, the divided cells did not repulse each other completely, but remained attached to each other to some extent in the two-cell and four-cell stages (Fig. 4A). From the observations, we reasoned that not only repulsion forces but also attraction forces might exist. We defined a parameter termed “stable repulsion ratio (*α*)” as the distance between the centers of the two cells (*d*_*ij*_) divided by the sum of radii of the two cells (*R*_*i*_*+ R*_*j*_) under the condition where the eggshell is removed (Fig. 4B). When there are only repulsion forces, *α* should be 1.0, because the two cells repulse each other until they are completely separated. For real embryonic cells, *α* was less than 1.0 (Fig. 4C). The results indicated that, when the distance between the centers of the cells is short (*d*_*ij*_ < *α*(*R*_*i*_ + *R*_*j*_)), the repulsion force is dominant, but when the distance increases (*d*_*ij*_ > *α*(*R*_*i*_ + *R*_*j*_)), the attraction force is dominant, so that the distance between the centers of the cells stabilizes at intermediate distance (*d*_*ij*_ = *α*(*R*_*i*_ + *R*_*j*_)).

**Figure 4.**
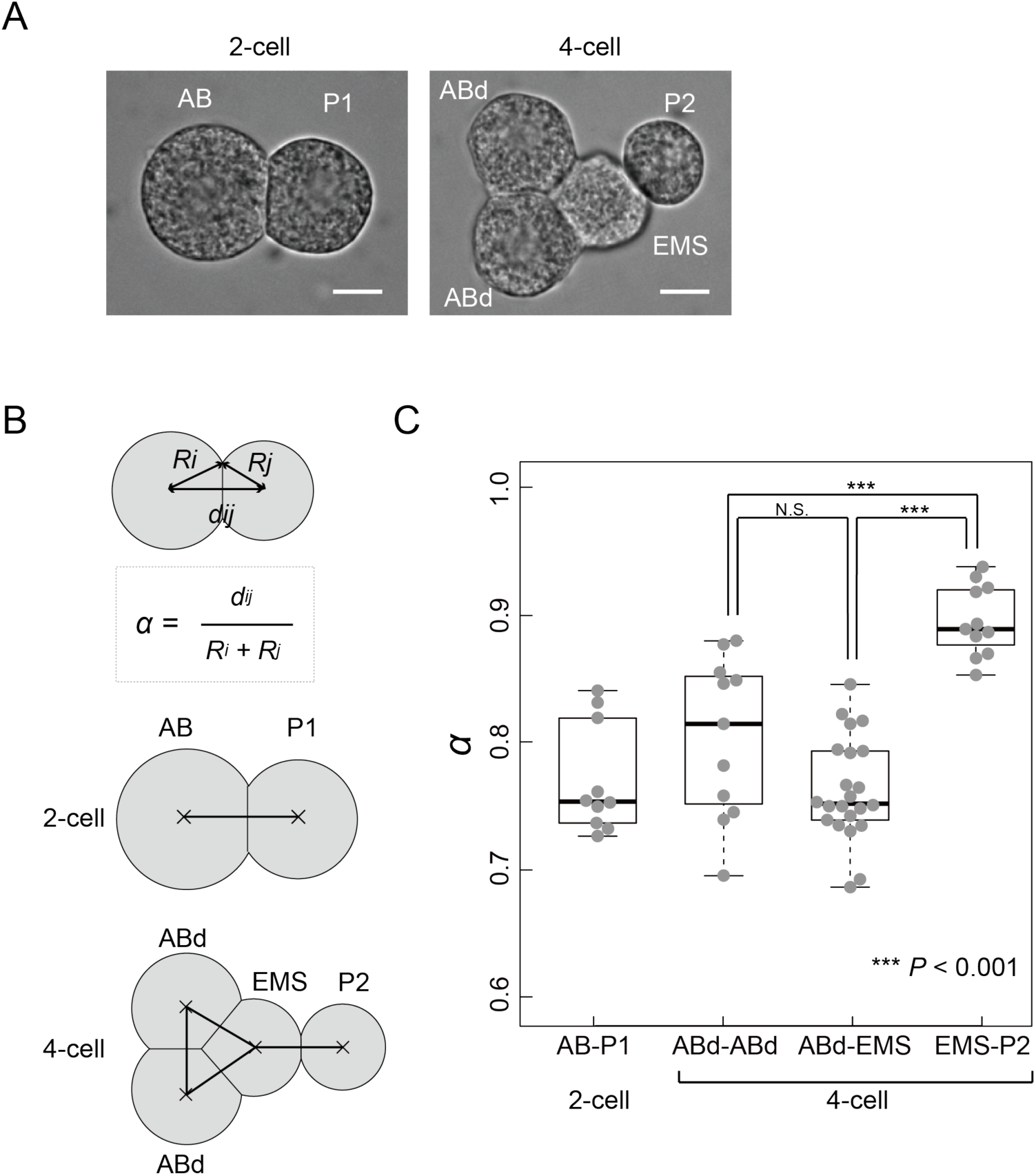
Anisotropic attraction forces among blastomeres at the four-cell stage in the *C. elegans* embryo. (A) Micrographs showing eggshell-removed *C. elegans* embryos at the two- or four-cell stage. The daughter cells of the AB cell were marked as ‘ABd‘ because we cannot distinguish ABa and ABp cells when the eggshell was removed. Scale bars are 10 μm. (B) Definition of stable repulsion ratio (*α*), which is calculated as the distance between cells (*d*_*ij*_) divided by the sum of radii (*R*_*i*_ + *R*_*j*_) of the cells. Diagram showing the combinations of the cells used to measure cell–cell distances at the two- or four-cell stage. (C) Bee swarm plot and boxplot of *α* in each combination of cell types (AB and P1 cells (two-cell stage), AB daughter cells (four-cell stage), AB daughter and EMS cells (four-cell stage), and EMS and P2 cells (four-cell stage)). \*\*\**P* < 0.001, Student’s *t*-test.

Moreover, we found that the degree of attraction force was not uniform but anisotropic, depending on the cell type. At the four-cell stage, ABa, ABp, and EMS cells were tightly attached (Fig. 4A), and *α* was approximately 0.75 (Fig. 4C). In contrast, P2 was loosely attached to EMS (Fig. 4A), and *α* for the EMS–P2 contact was approximately 0.90 (Fig. 4C). These results indicated that there are anisotropic attraction forces between the cells, in addition to the repulsion forces, at the four-cell stage of *C. elegans* embryos.

### Computer simulation of the anisotropic attraction (AA) model

To test whether the anisotropic attraction forces are the missing mechanism to explain the diversity and robustness of cell arrangements, we revised the RO model by adding the anisotropic attraction forces between the cells as observed *in vivo*. We termed the revised model as “the anisotropic attraction (AA) model.” In the AA model, we assumed that the intercellular force (*F*_*ij*_) becomes zero at intermediate distance when the distance between the centers of the two cells (*d*_*ij*_) is *α*(*R*_*i*_ + *R*_*j*_) (Fig. 5A). When *d*_*ij*_ > *α*(*R*_*i*_ + *R*_*j*_), the attraction force acts between the centers of the two cells as far as they are attached (*d*_*ij*_ < *R*_*i*_ *+ R*_*j*_) (Fig. 5A) and the distance between the cells gets closer until it reaches *d*_*ij*_ = *α*(*R*_*i*_ + *R*_*j*_). We set *α* to 0.90 for the interaction between EMS and P2 cells and to 0.75 for other interactions, based on the experimental measurements (Fig. 4C).

**Figure 5.**
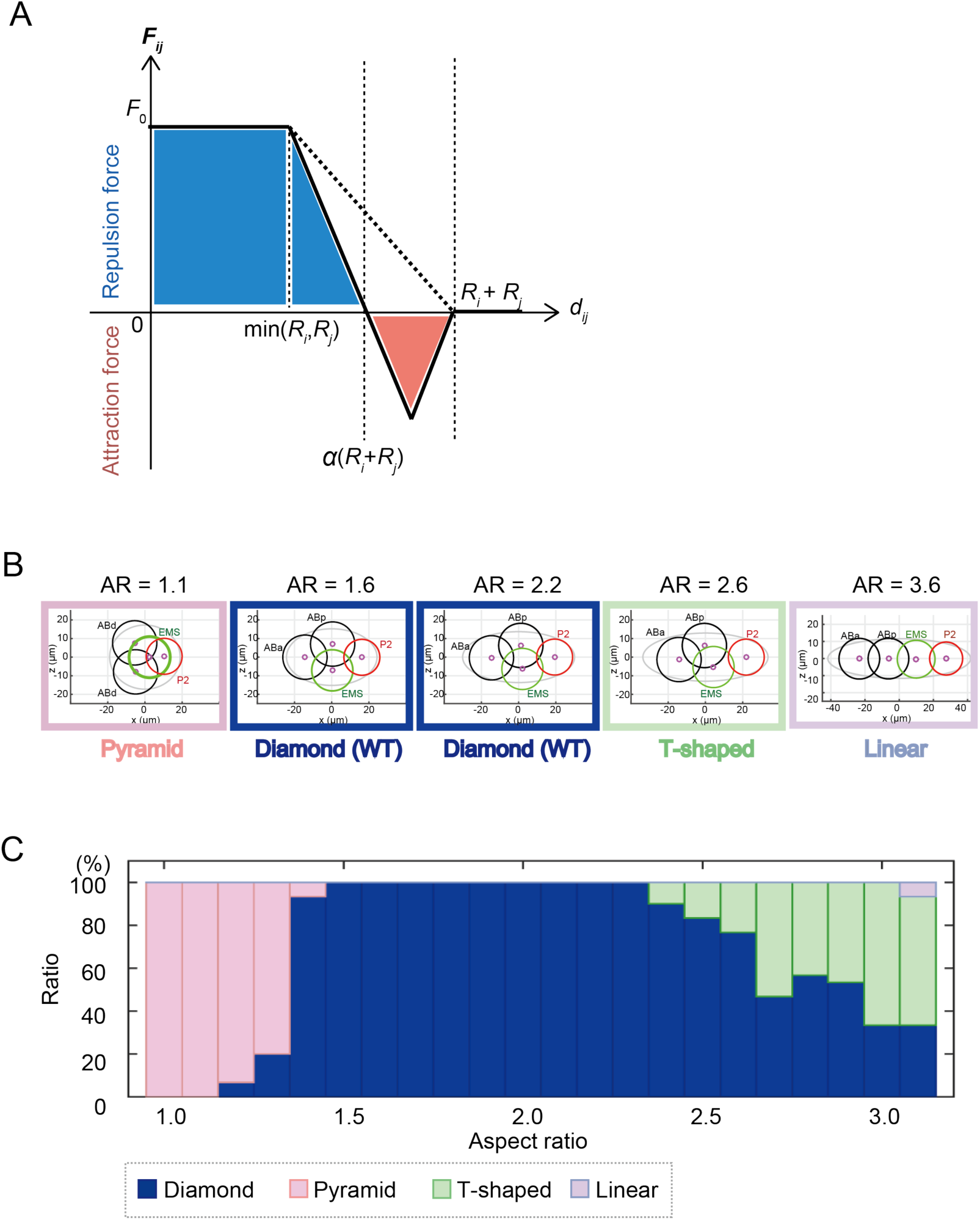
Simulation of cell arrangement patterns with the anisotropic attraction (AA) model. (A) Relationship between the strength of intercellular forces (*F*_*ij*_) and cell–cell distance (*d*_*ij*_) in the anisotropic attraction (AA) model. (B) Examples of the cell arrangement patterns at the four-cell stage simulated by the AA model at AR = 1.1, 1.6, 2.2, 2.6 and 3.6 respectively. (C) Relationship between rate of appearance of different types of cell arrangement (blue, diamond type; pink, pyramid type; green, T-shaped type; purple, linear type) and the ARs in a computer simulation based on the AA model (*α* = 0.90 (EMS and P2), and *α* = 0.75 (others)).

The behaviors predicted by the AA model resembled those of real embryos. Firstly, the AA model reproduced the diamond type of cell arrangement when the AR was 1.6 (Fig. 5B), which is the average AR for wild-type embryos measured in this study. Secondly, the model reproduced the pyramid-type arrangement when the AR was 1.1 (Fig. 5B). These two features were also observed in the RO model (Fig. 3C). Thirdly, and most importantly, the diamond-type arrangement was more robust against changes in the AR in the AA model than in the RO model. The AA model produced the diamond-type pattern even when the AR exceeded 3.0 (Fig. 5C), whereas the RO model did not produce the diamond-type pattern when the AR exceeded 2.0. Fourthly, the T-shaped type, which was observed *in vivo* for high ARs but was not reproduced in the RO model, was reproduced in the AA model (Fig. 5B, 5C). Fifthly, the AA model predicted the linear type for ARs exceeding 3.1 (Fig. 5B, 5C). In summary, the model taking into account the anisotropic attraction forces increased the robustness of the diamond-type cell arrangement and successfully reproduced all types of cell arrangement patterns observed in real embryos, including nematode species other than *C. elegans*. The modeling demonstrated that the framework of the AA model is sufficient to produce the diversity of cell arrangements and the robustness of the diamond type of cell arrangement.

### Attraction forces are produced by E-cadherin in the *C. elegans* embryo

Adhesion between cells can be explained by the difference in interfacial tensions between cells and between cells and the medium, as observed for contacting soap bubbles (Hayashi and Carthew, 2004; Pierre et al., 2016). Alternatively, cells may have an active mechanism to attract each other. Cell adhesion mediated by cadherins (Yoshida and Takeichi, 1982) provides such an active mechanism by localizing predominantly at the cell–cell contact region and attracting neighboring cells by opposing to surface tensions (Lecuit and Lenne, 2007). Among the cadherin genes, *hmr-1* encodes E-cadherin in *C. elegans*, and it has been shown that the HMR-1 protein localizes at cell adhesion sites from the embryonic stage onward (Grana et al., 2010; Pettitt, 2005). E-cadherin promotes the attraction between cells by increasing the area and strength of cell–cell contact, like a zipper (Lecuit and Lenne, 2007). In a transgenic *C. elegans* line expressing GFP fused to the HMR-1 protein under the control of the *mex-5* promoter that turns on gene expression maternally (Klompstra et al., 2015), we observed that the accumulation of HMR-1 at the cell boundaries was anisotropic and weak at the borders of the P2 cell (Fig 6A). The anisotropy was dependent on the *par-2* gene (Fig. 6A). The PAR-system, including PAR-2 protein, is a well-known pathway that maintains cell polarity in the *C. elegans* early embryo and in other species (Motegi and Seydoux, 2013; Munro et al., 2004). The localization of HMR-1 protein coincided with the anisotropy of the attraction forces observed *in vivo* (Fig. 4C).

**Figure 6.**
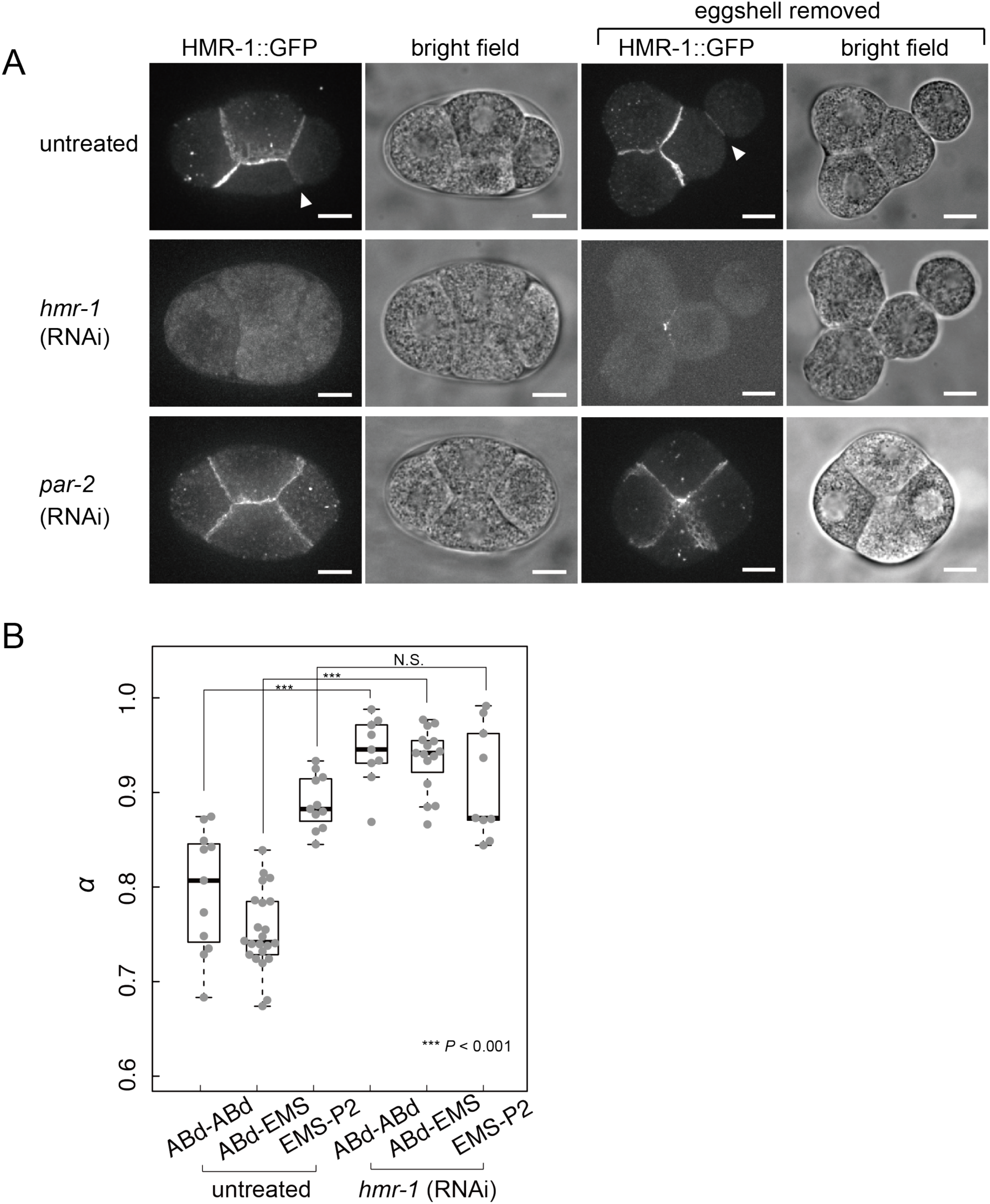
Cadherin localization and cell adhesion in the four-cell-stage *C. elegans* embryo. (A) Micrographs of embryos expressing HMR-1 fused with GFP protein in untreated, *hmr-1* RNAi- and *par-2* RNAi-treated embryos with or without eggshell. White arrowheads indicate cell–cell contact between EMS and P2 cell. Scale bars are 10 μm. (B) *α* of untreated and *hmr-1* RNAi-treated embryos without eggshell at the four-cell stage. \*\*\**P* < 0.001, Student’s *t*-test.

To test whether HMR-1 is required for the attraction forces, we knocked down *hmr-1* by RNAi. When the eggshell was removed, cell–cell adhesion was impaired, resulting in cells of nearly spherical shape (Fig. 6A). To quantify the degree of loss of adhesion, we measured the *α* parameter for *hmr-1* knockdown embryos. The parameter was significantly larger for *hmr-1* knockdown than for wild-type embryos (Fig. 6B). This indicated that the E-cadherin HMR-1 is necessary to produce the attraction forces at the four-cell stage. It should be noted that the attraction forces were not completely lost by the knockdown because the α parameter did not reach 1.0 even in the *hmr-1* RNAi condition (Fig. 6B).

### Attraction forces are necessary to produce the robustness of the diamond-type cell arrangement

From the physical modeling study (Fig. 5), we proposed that the attraction forces contribute to the robustness of the diamond-type arrangement against eggshell deformation. The model predicted that, without attraction forces, the diamond-type arrangement would be lost at high AR (Fig. 3D). To test this prediction, we attempted to experimentally reduce the attraction forces in embryos with high AR. In an initial attempt, we knocked down *hmr-1* alone by RNAi on the *lon-1*(*e185*) mutant background. However, the diamond-type arrangement was still observed in embryos with high ARs over 2.0 (Fig. S4A). We suspected that the attraction forces were not completely lost under *hmr-1* RNAi, which is consistent with the fact that the *α* parameter did not reach 1.0 in this condition (Fig. 6B).

We next simultaneously knocked down *hmr-1* (E-cadherin) and *hmp-2* (β-catenin) by RNAi. β-Catenin mediates forces downstream of the adhesion by E-cadherin to cortical actin networks, and this downstream signaling is suggested to activate cell adhesion (Lecuit and Lenne, 2007) (Fig. 7A). Therefore, we expected that the knockdown of β-catenin would further impair the attraction forces between cells. Indeed, the majority of *hmr-1; hmp-2* knockdown embryos lost the diamond-type arrangement at ARs over 2.0 (Fig. 7B, S4A). *hmr-1; hmp-2* knockdown embryos were able to take the diamond-type arrangement in the normal range of ARs (1.6–2.0) (Fig. S4B), indicating that the robustness of the diamond-type arrangement was impaired by the knockdown. The experimental results agreed with the prediction from the model without attraction forces, (i.e., the RO model), which predicted that, without attraction forces, the range of ARs to support the diamond-type was narrower and thus, less robust (Fig. 3D). It should be noted that the model simulation predicted that the linear type of cell arrangement would appear; however, the majority of real embryos adopted a T-reverse type at high ARs (Fig. 7B), in which ABp was in contact with EMS and P2, while ABa and EMS did not contact (Fig. 7C) (see Discussion for possible explanations). Overall, we demonstrated that the attraction forces via cell adhesion produce the robustness of the proper cell arrangement against eggshell deformation both theoretically and experimentally.

**Figure 7.**
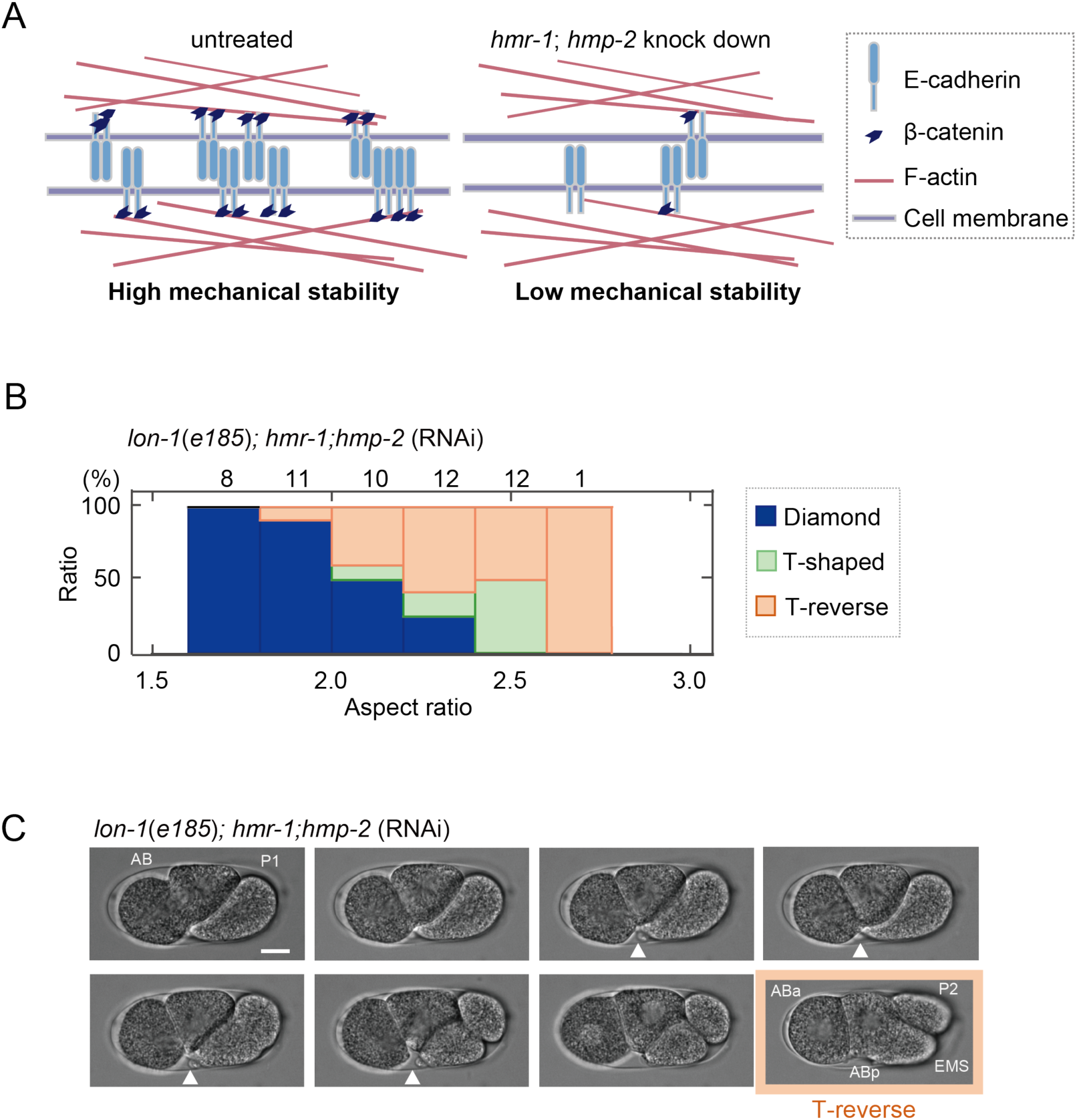
Impaired robustness of the diamond-type arrangement against eggshell deformation in the *C. elegans* embryo. (A) Model of E-cadherin and β-catenin-mediated high mechanical stability of the cell arrangement pattern through interactions with the cortical actin network. (B) Relationship between the rate of appearance of the patterns of cell arrangements (blue, diamond type; green, T-shaped type; orange, T-reverse type) and the ARs in the *hmr-1*; *hmp-2* knockdown strain with *lon-1(e185)* background. The numbers above the bars represent the number of embryos. (C) Sequential snapshots when breakage of cell adhesion between ABa and EMS cells occurred in the *hmr-1; hmp-2* double knockdown strain with *lon-1*(*e185*) mutant background. White arrowheads indicate cell–cell contact between ABa and future EMS cell. T-reverse-type cell arrangement (absence of cell–cell contact between ABa and P2 cells, and ABa and EMS cells) was formed at the four-cell stage (orange). Scale bar is 10 μm.

## Discussion

### Diverse patterns of cell arrangements in nematode embryos in the four-cell stage

The patterns of cell arrangements in the four-cell stage of various nematode species can be classified into four different types. The patterns correlated with the shapes (i.e., the AR) of the eggshells. In this study, we succeeded in changing the diamond-type cell arrangement of *C. elegans* embryos into the other three types, demonstrating that the eggshell shape is sufficient to change the cell arrangement pattern. The aspect-ratio-dependency of these patterns in various nematode species resembled that in *C. elegans*. Because our AA model accounts for the AR-dependency in *C. elegans*, it well explains the diversity in cell arrangement patterns in various nematode species.

The AR-dependency of the patterns of cell arrangements in various nematode species was not exactly the same as that in *C. elegans*. An apparent difference is that some nematode species with ARs of 2.0–2.6 take a linear-type arrangement (Fig. 1C) whereas *C. elegans* embryos with ARs in this range do not (Fig. 2C). The arrangements in these species can be explained with the AA model. In the absence of attraction forces, the AA model coincides with the RO model (Fickentscher et al., 2013) by definition. The RO model predicts the linear-type arrangement when the AR exceeds 2.0 (Fig. 3D). Therefore, the nematode species that take the linear type with ARs of 2.0–2.6 are explained by the AA model with reduced attraction forces.

The patterns of cell arrangement are determined by the coordination of cell division orientation, position, and timing in addition to physical interaction among cells and the spatial confinement of the eggshell. Our model is capable to examine effects of changing those parameters. For example, the orientation of cell division is a parameter in our AA model, and thus, we can change the orientation in the model and compute the consequences. *par-2* and *par-3* mutants are well known for their change in the orientation of cell division in the two-cell stage *C. elegans* embryo. To test whether our model accounts for the mutants’ arrangement, however, we need to change not only the orientation but also the attraction forces, as the localization of cadherins is also affected by the mutations (Munro et al., 2004). Another complexity is that cell shapes affect the orientation of cell division axis. The mitotic spindle tends to align along the long axis of the cell and the cell division plane forms perpendicular to the spindle, known as Hertwig’s rule (Kimura and Onami, 2007; Minc et al., 2011; Pierre et al., 2016; Dumollard et al., 2017). The cell arrangement is thus determined by the interplay of orientation of cell division, cell shape, and eggshell, as well as the strength of attraction and repulsion forces. Our AA model will be a good framework to integrate these parameters in one model.

### Changing eggshell shape through gene manipulation

In this study, we obtained eggshells with various ARs by using gene manipulations. We focused on the genes that affect the thickness of the adult body, more specifically, the thickness of the gonad. We expected the eggs that passed through the thicker gonad would be rounder (Fig. S1C). Indeed, the *dpy-11* and *lon-1* mutants, whose adult bodies are thick and thin, respectively, produced round and slender eggs, respectively. In addition, by increasing the volume of the egg by *C27D9.1* RNAi, we were able to obtain slender eggs possibly because the relative thickness of the gonad is thinner (Fig. S1). It would be an interesting future challenge to investigate the correlation between the thickness of the gonad and the shape of the eggs through examining the shapes of various nematode species (Goldstein, 2001; Schulze and Schierenberg, 2011).

### Roles of E-cadherin and β-catenin in the attraction forces

We showed that the attraction forces between cells are important for the diversity and robustness of cell arrangements. We found that the attraction forces were impaired by knocking down *hmr-1*/E-cadherin. By knocking down both *hmr-1*/E-cadherin and *hmp-2*/β-catenin, the robustness of the diamond-type arrangement was impaired. From the results, we concluded that E-cadherin and β-catenin are responsible for generating attraction forces. In our measurement of the α parameter in embryos without eggshell (Fig. 6), the knockdown of *hmr-1*/E-cadherin alone severely, but not completely, impaired the attraction forces. We had to knockdown *hmp-2* in addition to *hmr-1* to observe the loss of robustness in embryos inside the eggshell. One possibility for this is that *hmr-1; hmp-2* RNAi caused an uncharacterized defect independent of the defect in attraction forces that affected the robustness of the diamond-type arrangement. We favor, however, a scenario involving the attraction forces because once attached, ABa and EMS cells separated in *hmr-1*; *hmp-2* knockdown embryos with eggshells with high ARs. We speculate that pressures provided by the spatial constraints of the eggshell increased the attachment between the cells and bypassed the requirement for E-cadherin to some extent. The effective attraction forces in *hmr-1* RNAi embryos within the eggshell might be larger than those in embryos without eggshell. This scenario explains why the knockdown of *hmr-1* alone was insufficient.

### Limitations of the AA model

One apparent discrepancy between the cell arrangements in the *C. elegans* embryos and our model is that, without the attraction forces in the AA model (i.e., RO model), the linear type of cell arrangement was expected for longer embryos, while real embryos took a T-reverse-type arrangement in *hmr-1*; *hmp-2* RNAi (Fig. 3D, 7B, 7C). There are two possible reasons for the discrepancy. Firstly, the cell shape might determine the orientation of the cell division axis. In slender eggshells, cells elongate along the long axis at the two-cell stage. The cell division axis of the elongated AB cell might tilt toward the long axis, because it tends to divide along the long axis of the cell (Hertwig’s rule) (Minc et al., 2011). Secondly, while the AA model assumed the eggshell to be ellipsoidal, real eggshells with high ARs were thicker near the poles. EMS and P2 cells can arrange perpendicular to the long axis in the thick region, with enough space to position cells according to a T-reverse type of cell arrangement.

### Significance of the attraction forces in embryo organization

The importance of the attraction forces is expected not to be limited to the four-cell stage of *C. elegans* embryos, but to be general. E-cadherin and β-catenin are generally localized at cell–cell contacts in embryos and tissues in different stages and in different animals. In early mouse embryos, the area of cell–cell contact increases and the cell shape becomes flat in an E-cadherin-dependent manner during the compaction process (Larue et al., 1994). The compaction process can be explained by an increase in attraction forces by E-cadherin upregulation in specific cells. In contrast, downregulation of attraction forces may be important in other cases. In sea urchin embryos, β-catenin signal on the cell membrane decreases in the primary mesenchyme cells, which leads to their detachment from neighboring cells and shift into blastocoel during gastrulation (Miller and McClay, 1997). Our present study added another important role of attraction forces between the cells to produce diversity and robustness of cell arrangement.

### Materials and methods

#### *C. elegans* strains

The *C. elegans* strains used in this study are listed in Table S2. The *C. elegans* strains and *Diploscapter coronata* were maintained using a standard *C. elegans* procedure (Brenner, 1974; Stiernagle, 2006). *Aphelenchoides besseyi* was maintained on PDA plates cultured with *Botryllus cinera* (Yoshida et al., 2009).

#### RNAi

Genetic knockdown of *C27D9.1*, *par-2*, *hmr-1*, and *hmp-2* was established by feeding RNAi as described previously (Kamath et al., 2000). For *hmr-1* RNAi, a probe targeting base pairs 13,610 to 14,254 of the unspliced *hmr-1* gene was used. For other genes, the *C. elegans* RNAi library (Source BioScience, Nottingham, UK) was used (Fraser et al., 2000).

#### Eggshell removal

Eggshells were removed using a modified previously described method (Park and Priess, 2003). Embryos were treated with bleaching agent (Kao, Japan), mixed with 10 N KOH in a 3:1 ratio for 90 s, and placed in Shelton’s growth medium (Shelton and Bowerman, 1996) for washing three times. The vitelline membrane was removed by trituration using a 30-μm micropipette made by pulling a glass capillary (GD-1; Narishige, Japan) with a micropipette puller (P-1000IVF; Sutter Instrument, Novato, CA, USA).

#### Image acquisition

Embryos were placed in 75% egg salt (or in Shelton’s growth medium in case the eggshell was removed). For fluorescence images, embryos were visualized at room temperature (22–24°C) using a spinning-disk confocal system (CSU-X1; Yokogawa Electric Corporation, Japan) mounted on an inverted microscope (IX71; Olympus, Japan) equipped with a 60×, 1.30 N.A. objective (UPLSAPO 60XSO; Olympus). Digital images were acquired with a CCD camera (iXon; Andor Technology, Belfast, UK) controlled by Metamorph software (version 7.7.10.0). Images are shown as maximum-intensity projections of planes spaced 1.0 μm apart. Images were analyzed with ImageJ (National Institute of Health, Bethesda, MS, USA).

Phase contrast images were acquired at room temperature (22–24°C) under an inverted microscope (Axiovert 100; Carl Zeiss, Germany) equipped with a 40×, 0.70 N.A. objective (Plan-Neofluar; Carl Zeiss). Digital images were acquired with a CCD camera (ORCA-100; Hamamatsu, Japan) controlled by iVision-Mac (version 4.0.9; BioVision Technologies, Exton, PA, USA). The AR and *α* were quantified with ImageJ. For the quantification of *α,* each blastomere was fitted into a precise circle by hand, and *R*_*i*_ and *d*_*ij*_ were quantified. The parameter *d*_*ij*_ was calculated from the centroids of the cells.

#### Statistical analysis

To confirm normality, the Shapiro–Wilk test was used. To confirm homoscedasticity, an F-test was used. If both normality and homoscedasticity were confirmed, Student’s *t*-test was used to compare means, and if only normality was confirmed, Welch’s *t*-test was used. In other cases, Wilcoxon’s rank sum test was used to compare mean values. *P*-values < 0.05 were considered statistically significant. For these analyses, R (<u>www.r-project.org</u>) was used. The experiments were not randomized, and the investigators were not blinded to allocation during experiments and outcome assessment.

#### Construction of the AA model based on RO model, and computer simulations

Three-dimensional simulations of cell motions inside the confined space (eggshell) were constructed by modifying the simulation developed by Dr. Weiss and colleagues (Fickentscher et al., 2013). The mathematical models consider cells to be soft spherical balls and the eggshell as a rigid ellipsoid. From an initial configuration, configurations of cells were calculated at successive time steps. We added the attraction forces in this study. We changed the AR value from 1.0 to 4.0 using 0.1 increments, while maintaining the total volume of the eggshell constant. The parameters used are listed in Table S3. The simulations were programmed in MATLAB and source code is available upon request.

The eggshell was considered to be an ellipsoid of which the center is the origin and the long axis is on the x axis in coordinate, with the length of the long axis defined as *lx,* and the length of short axes as *ly* (= *lz*). The AR is calculated as *lx*/*ly*. For the simulation, the AR value was changed from 1.0 to 4.0 using 0.1 increments, while maintaining the total volume of the eggshell constant (Table S3). Cells were considered to be spheres of radius *R*_*i*_ (*i* = 1, 2,…, *N*, with *N* denoting the total number of cells). The center of mass is represented by position **r**_i_.

Cells were assumed to move in a highly viscous environment, and the positions were calculated by an overdamped Langevin’s equation as follows:

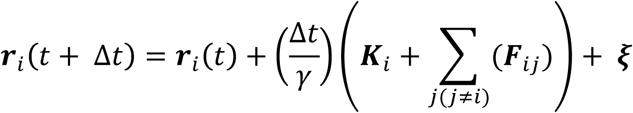

Random cell motion due to stochastic effects is represented by a vector ***ξ***. The components are three combinations of uncorrelated random numbers with a mean of 0 and variance of 0.027. The integration time step *Δt = 5* s.

##### Repulsion forces from the eggshell

When touching the eggshell, the cells experience a repulsion force defined as

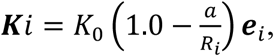

where ***e***_*i*_ denotes a unit vector perpendicular to the eggshell, pointing into the cells, and *a* is the minimum distance between the center of cell *i* and the eggshell. *a* was calculated as the minimum solution of the following equation that calculates the distances from a coordinate inside the ellipsoid to the nearest edge of the ellipsoid. The equation is derived when we ask the components of the vector perpendicular to the tangent plane on the ellipsoid.

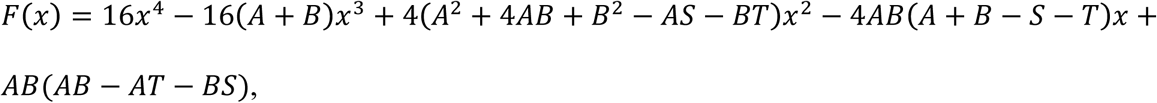

where *A* denotes the square of *lx*, *B* denotes the square of *ly*, *S* denotes the square of the x coordinate of the cell, *T* denotes the sum of squares of the y and z coordinates of the cell. *a* was calculated by the function equipped in MATLAB. For *a* > *R*_i_, *K*_i_ = 0.

##### Repulsion and attraction forces between cells

The force between a pair of cells depends on the distance between the centers of the cells (cell *i* and cell *j*), *d*_*ij*_ = |***r***_*i*_ *-* ***r***_*j*_|. For *d*_*ij*_ < *α*(*R*_*i*_ + *R*_*j*_), two cells repulse each other, and for *α*(*R*_*i*_ + *R*_*j*_) < *d*_*ij*_ < (*R*_*i*_ + *R*_*j*_), two cells attract each other. Otherwise, the pairwise force is zero. Specifically, forces between any two cells were calculated as follows.

For 0 < *d*_*ij*_ < min(*R*_*i*_, *R*_*j*_), **F_*ij*_** = *F*_0_**e_*ij*_**.

For min(R_i_, R_j_) < *d*_*ij*_ < 0.5 (1+α) (R_i_+R_j_), 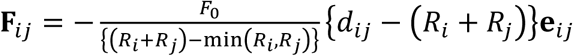.

For 0.5 (1+α) (R_i_+R_j_)< *d*_*ij*_ < (R_i_+R_j_), 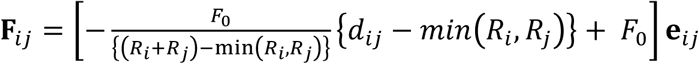.

Otherwise, **F**_*ij*_ = **0**.

Here, **e**_*ij*_ *=* **d**_*ij*_ */ |***d**_*ij*_*|* is the unit vector pointing from cell *i* to cell *j*. *R*_*i*_ and *R*_*j*_ is the radii of cell *i* and cell *j*, respectively. *α* is the parameter at which the pairwise force is zero between two attached cells. In the expressions, min() represents the minimum value of the parameter in parentheses. The constant force for *0 < d*_*ij*_ *<* min(*R*_*i*_*, R*_*j*_) reflects cell elongation during cytokinesis, which occurs roughly at a constant velocity.

##### Cell divisions

The orientations of the division axes of the first (P0 to AB and P1), second (AB to ABa and ABp), and third division (P1 to EMS and P2) are parallel to the X-, Z-, and X-axis, respectively.

##### Initial configuration

At the initial time step, two cells are positioned at the center of the eggshell 2.0 μm apart on the X-axis because the first cell division orientation is parallel to the long axis of the embryo. The patterns of cell arrangements were classified based on whether the cell–cell distance (*d*_*ij*_) is smaller or larger than the sum of radii (*R*_*i*_ + *R*_*j*_) in each combination of two cells. When the cell–cell distance is smaller than the sum of radii, we considered that the cells contacted each other.

## Acknowledgements

We thank Drs. Matthias Weiss and Rolf Fickentscher (University of Bayreuth) for providing the source code of their model, Kenji Sugioka (University of Oregon) for technical advice, Shuichi Onami (RIKEN), Hitoshi Sawa, Yumiko Saga, Daiju Kitagawa, Yoshihisa Oda, Yuta Shimamoto, Jun Takagi (National Institute of Genetics), Akira Funahashi, Mikiko Motomuro (Keio University) and the members of our lab for discussion. Some of the strains were provided by the *Caenorhabditis* Genetics Center, funded by the National Institute of Health. Drs. Yuji Kohara, Hiroshi Kagoshima (National Institute of Genetics), and Toyoji Yoshiga (Saga University) provided us with *Diploscapter coronata* and *Aphelenchoides besseyi*. This project was supported by a JSPS doctoral fellowship (JP16J09469) to K.Y., JSPS KAKENHI grant numbers JP15H04372, JP15KT0083 to A.K., and by the Naito Foundation and the Sumitomo Foundation.

## Author contributions

K.Y. and A.K. conceived the study, analyzed the data and wrote the manuscript. K.Y. performed all the experiments, constructed the models, and ran the calculations.

